# PlantRNA-FM: An Interpretable RNA Foundation Model for Exploration Functional RNA Motifs in Plants

**DOI:** 10.1101/2024.06.24.600509

**Authors:** Haopeng Yu, Heng Yang, Wenqing Sun, Zongyun Yan, Xiaofei Yang, Huakun Zhang, Yiliang Ding, Ke Li

**Author notes:** To whom correspondence should be addressed. Correspondence may also be addressed to or. The authors wish it to be known that, in their opinion, the first four authors should be regarded as Joint First Authors.

## Abstract

The complex ‘language’ of plant RNA encodes a vast array of biological regulatory elements that orchestrate crucial aspects of plant growth, development, and adaptation to environmental stresses. Recent advancements in foundation models (FMs) have demonstrated their unprecedented potential to decipher complex ‘language’ in biology. In this study, we introduced PlantRNA-FM, a novel high-performance and interpretable RNA FM specifically designed based on RNA features including both sequence and structure. PlantRNA-FM was pre-trained on an extensive dataset, integrating RNA sequences and RNA structure information from 1,124 distinct plant species. PlantRNA-FM exhibits superior performance in plant-specific downstream tasks, such as plant RNA annotation prediction and RNA translation efficiency (TE) prediction. Compared to the second-best FMs, PlantRNA-FM achieved an *F*1 score improvement of up to 52.45% in RNA genic region annotation prediction and up to 15.30% in translation efficiency prediction, respectively. Our PlantRNA-FM is empowered by our interpretable framework that facilitates the identification of biologically functional RNA sequence and structure motifs, including both RNA secondary and tertiary structure motifs across transcriptomes. Through experimental validations, we revealed novel translation-associated RNA motifs in plants. Our PlantRNA-FM also highlighted the importance of the position information of these functional RNA motifs in genic regions. Taken together, our PlantRNA-FM facilitates the exploration of functional RNA motifs across the complexity of transcriptomes, empowering plant scientists with novel capabilities for programming RNA codes in plants.

## Introduction

The transcriptome contains a wide array of RNA motifs that impact diverse biological functions such as translation^1–5^. These RNA motifs encompass both RNA sequence and structure features. Previous individual studies have revealed the functional importance of RNA sequence features such as the Kozak sequence motif^6^. Recently, our studies along with others have suggested that both RNA secondary and tertiary structure motifs play important roles in diverse biological processes^7–13^. Particularly in plants, the relatively low habitat temperatures (∼20 °C) favour the folding of RNA structure motifs, including RNA tertiary motifs such as RNA G-quadruplex (rG4)^12^. However, systematically identifying functional RNA motifs across transcriptomes remains a formidable challenge due to the high level of complexity arising from astronomical combinations of the four nucleotide bases into tens of thousands of transcripts^8,14^. For example, for a 50-nucleotide sequence, the number of artificially synthesized sequences would be on the order of 4^50^ (approximately 1.27 × 10^30^), which is impossible to achieve experimentally. Additionally, the functional readouts using the reporter gene assay for measuring biological functions such as translation may not be sensitive enough to detect differences in individual single-nucleotide mutations^15^.

The recent rapid advancements of foundation models (FMs) in artificial intelligence (AI) are set to show exciting promise for supercharging scientific advances in life sciences^16^. FMs are distinguished by their massive scale, often encompassing millions to billions of parameters. They are first pre-trained in a self-supervised manner on diverse forms of unlabelled data. This makes them ideally suitable for bioscience, where acquiring abundant labelled data is both prohibitively expensive and time-consuming. More importantly, FMs are highly adaptable through fine-tuning and are poised to aid bioscientists in customising generalist FMs in unravelling complex biological processes, paving the way for unprecedented capabilities in modulating gene functions. For FMs on DNA sequences, DNABERT2 is one of the FMs pre-trained on the genome sequences across 135 species, including mammals, fungi and bacteria^17^. By pre-training on diverse human and non-human genomes, the Nucleotide Transformers (NT) family learns transferable representations that enable accurate molecular phenotype prediction with limited annotated data, while focusing on key genomic elements without supervision^18^. FMs have also achieved success in protein sequences, also known as protein language models. For example, ESM2 (Evolutionary Scale Modeling) has achieved remarkable breakthroughs in atomic-level structure representations by pretraining on a vast amount of protein sequences and structures^19^.

For building RNA FM, several FM models were pre-trained using RNA sequence information that has demonstrated great performance in RNA molecule design^20–22^. However, RNA sequence information is not sufficient since RNA is capable of forming secondary or tertiary structure motifs that are important for its functions^23,24^. Therefore, it is important to generate an RNA FM including both RNA sequence and structure information to facilitate the exploration of functional RNA motifs. Here, we developed PlantRNA-FM, a groundbreaking RNA FM designed to globally identify functional RNA motifs including both RNA sequences and structure motifs in plants (Fig. 1). By incorporating RNA sequences, annotations, and structure information from 1,124 distinct plant species, PlantRNA-FM captures the extensive diversity of plant transcriptomes (Fig. 1). We validate the superior performance of PlantRNA-FM in downstream tasks compared to existing FMs. Furthermore, we also established an interpretable framework based on our PlantRNA-FM to determine the critical regions across the 5’ untranslated regions (5’ UTRs) that significantly impact translation. Remarkably, PlantRNA-FM identifies RNA motifs at the transcriptome-wide scale that are functionally important to translation including both RNA sequences, and secondary and tertiary structure motifs. We further experimentally validated these identified RNA motifs in plants. The development of our PlantRNA-FM represents a significant leap forward in our ability to decipher hidden regulatory codes among the extensive complexity of nucleotides across the transcriptome, opening new avenues for RNA-based gene regulation.

**Fig. 1.**
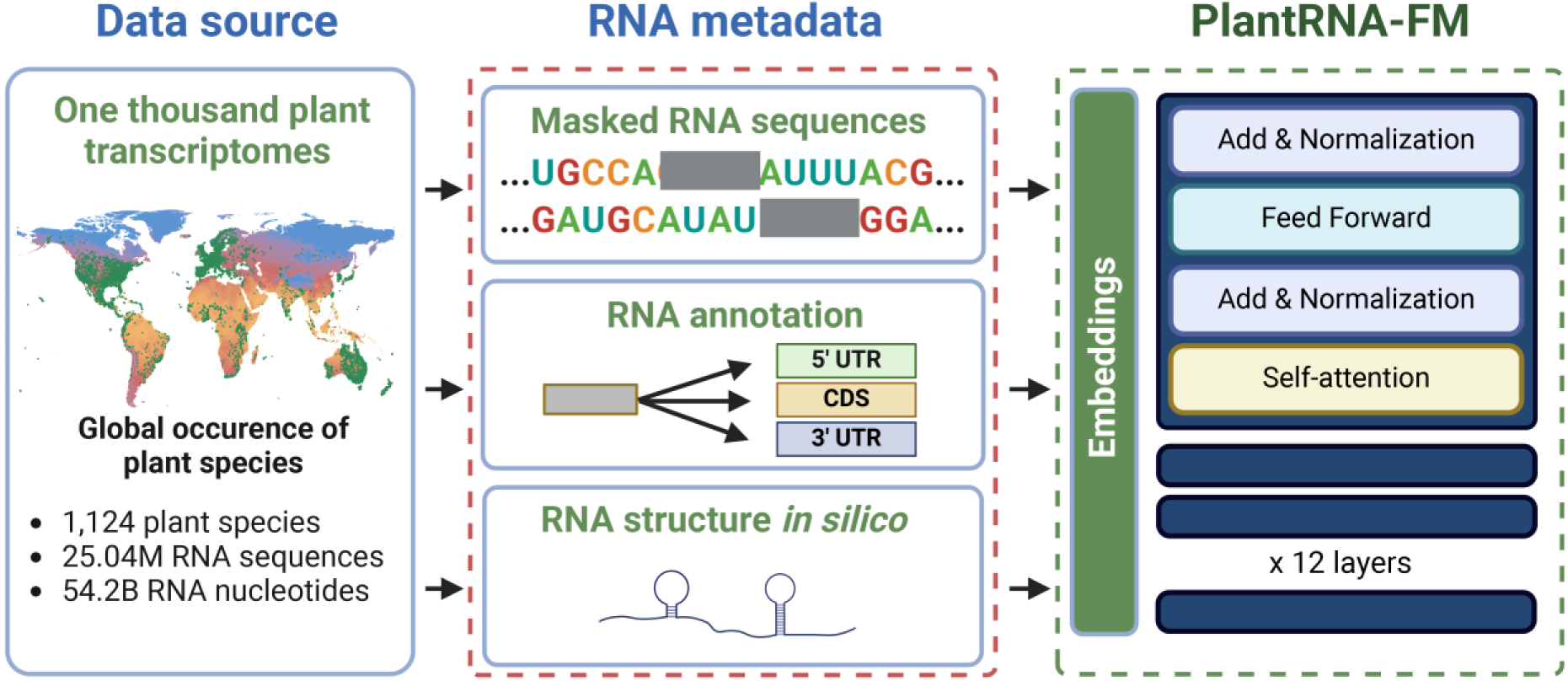
Schematic overview of the Pre-training Phase of PlantRNA-FM. The pre-training dataset comprises transcriptomic sequences from 1,124 plant species, consisting of approximately 25.0M RNA sequences and 54.2B RNA bases. The green dots on the global mean temperature map represent the geographical distribution of these plant species across the world.

## 1. Results

### 1.1. Our PlantRNA-FM integrates both RNA sequence and structure information of the transcriptomes across 1,124 plant species

The plant kingdom encompasses approximately 500,000 species, exhibiting remarkable diversity. The One Thousand Plant Transcriptomes Initiative (1KP) sequenced the transcriptomes of 1,124 species, capturing the extensive diversity of plant transcriptomes^14^. Here, we took advantage of this unique resource and generated the pre-training dataset for our PlantRNA-FM (Fig. 1). Different from existing FMs, our PlantRNA-FM was designed to capture and learn both RNA sequences and RNA structure motifs. We employed *RNAfold*^25^ to predict RNA structures of individual RNA sequences across 1,124 transcriptomes and integrated them into the pre-training dataset. Our PlantRNA-FM has 35 million parameters, including 12 transformer network layers, 24 attention heads, and an embedding dimension of 480, optimised for RNA understanding rather than generation (**Methods**). Our tokenization approach surpasses the constraints of conventional *k*-mers and BPE methods, ensuring the preservation of RNA structure motifs as coherent units throughout the pre-training process (**Methods**). In addition, we incorporated RNA annotation information (CDS and UTRs) and employed advanced pre-training techniques, such as sequence truncation, filtering and masked nucleotide modeling (**Methods**).

To assess the effectiveness of our PlantRNA-FM in RNA structure prediction tasks, we evaluated its performance (Fig. S1, Table S1) using three benchmark datasets: bpRNA, ArchiveII, and RNAstralign^26–28^. The *F*1 score, which is the harmonic mean of precision and recall, was used to measure the model’s predictive performance on these datasets. The *F*1 scores achieved by our PlantRNA-FM on these three datasets were 0.750, 0.924, and 0.981, respectively, while *RNAfold* alone only obtained *F*1 scores of 0.278, 0.759, and 0.748 (Fig. S1, Table S1). When compared to other state-of-the-art FMs, PlantRNA-FM outperformed the second-best model by 22.10%, 27.49%, and 17.38% on the respective datasets (Fig. S1, Table S1). Therefore, the unique integration of RNA structure information equips our PlantRNA-FM with the ability to predict RNA structure more accurately.

### 1.2. PlantRNA-FM demonstrates superior performance on plant-specific downstream tasks

To evaluate the performance of PlantRNA-FM, we curated a benchmark set consisting of four other state-of-the-art FMs: DNABERT-2, Nucleotide Transformer, ESM2, and cdsBERT. We assessed their performance in two plant-specific downstream tasks: genic region annotation and translation efficiency (TE) prediction (Fig. 2a).

**Fig. 2.**
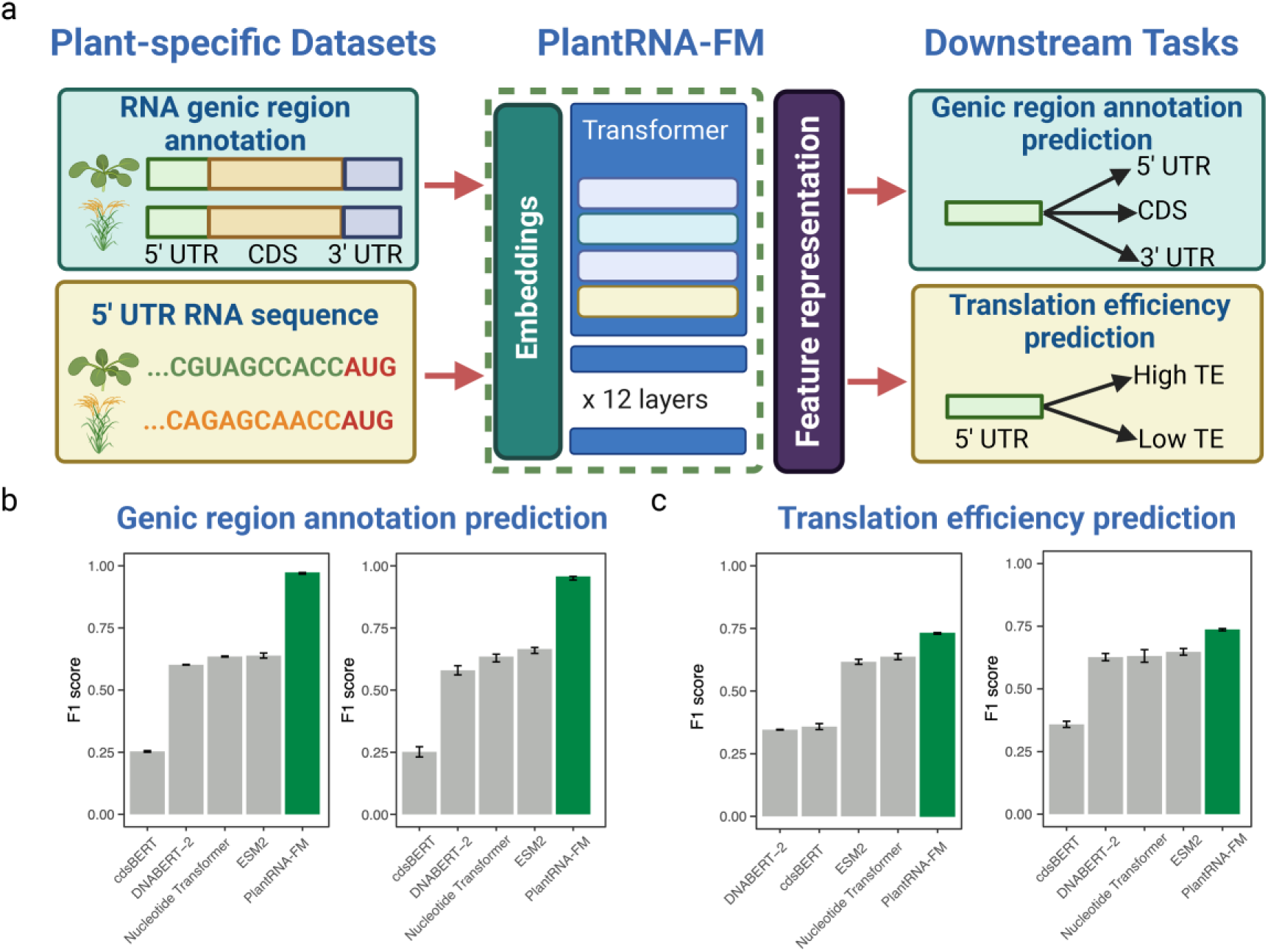
Fine-tuning PlantRNA-FM on plant-specific datasets. a, Overview of fine-tuning PlantRNA-FM for RNA genic region annotation prediction and RNA translation efficiency (TE) prediction tasks. *A. thaliana* and *O. sativa* were selected as representative plant species. For the RNA genic region annotation prediction task, RNA sequences from these two species were included, along with three labels: 5’ UTR, CDS, and 3’ UTR. For the RNA TE prediction task, 5’UTR sequences from these two species were included, along with TE labels (high TE and low TE). b, c, Comparison of the model performance of different pre-trained models on RNA genic region annotation prediction and RNA TE prediction tasks. The error bars represent the standard deviation of the *F1* scores obtained from three fine-tuning replicates.

In the RNA genic region annotation prediction task, we aimed to identify and classify different genic regions of given RNA sequences, such as the 5’ UTR, coding sequence (CDS), and 3’ UTR. We used the transcriptomes of two model plant species, *Arabidopsis thaliana* (a dicot model plant) and *Oryza sativa L.* ssp*. Japonica* (rice, a moncot model plant). Both of them were not included in our pre-training dataset. For the RNA genic region annotation prediction in these two species, our PlantRNA-FM outperformed other FM models, achieving average *F*1 scores of 0.974 and 0.958 for *Arabidopsis* and rice, respectively, surpassing the second-best model by 52.45% and 43.90% (Fig. 2b, Table 1).

**Table 1.** Comparison of *F1* scores achieved by different pre-trained models on benchmark datasets.

| Tasks | Species | PlantRNA-FM | cdsBERT | DNABERT-2 | Nucleotide Transformer | ESM2 |
| --- | --- | --- | --- | --- | --- | --- |
| RNA genic region annotation prediction | *A. thaliana* | 0.974±0.003 | 0.254±0.003 | 0.602±0.001 | 0.635±0.002 | 0.639±0.008 |
|  | *O. sativa* | 0.958±0.006 | 0.252±0.017 | 0.580±0.015 | 0.635±0.013 | 0.665±0.010 |
| RNA translation efficiency prediction | *A. thaliana* | 0.735±0.003 | 0.359±0.010 | 0.346±0.001 | 0.637±0.010 | 0.617±0.008 |
|  | *O. sativa* | 0.737±0.004 | 0.359±0.010 | 0.627±0.012 | 0.631±0.020 | 0.649±0.011 |

For translation, one of the key RNA biological processes, previous research has highlighted the critical role of the 5’ UTR in regulating translation efficiencies^17–19,21,29–31^. To evaluate the TE prediction performance of our PlantRNA-FM, we used the 5’ UTR sequences of both *Arabidopsis* and rice transcriptomes along with the corresponding TE values measured by polysome profiling^8^. We first classified the TE datasets into high and low TE groups, using the mean plus or minus the standard deviation as the threshold. In the TE prediction task, PlantRNA-FM achieved *F*1 scores of 0.735 and 0.737 for *Arabidopsis* and rice, respectively, outperforming the second-best model by 15.30% and 13.83% (Fig. 2c). Taken together, our PlantRNA-FM is better suited for plant-specific downstream tasks compared to other FMs pre-trained on non-plant datasets.

### 1.3. Interpretable PlantRNA-FM revealed RNA features important to translation

A general roadblock in applying AI models to biology is that, while these models demonstrate strong predictive capabilities, the key to their successful application lies in interpreting them to uncover the biological principles learned by the AI. In this paper, we established an interpretable framework to derive an attention contrast matrix from our PlantRNA-FM (**Methods**). In particular, we are interested in extracting the key RNA features within the 5’ UTR that significantly impact RNA translation, i.e., elucidating the RNA motifs associated with translation (Fig. 3a). We developed two models in parallel: one is the true model, denoted as PlantRNA-FM(+), trained using the real TE dataset, while the other one is called the background model, PlantRNA-FM(-), altered using the same dataset but with randomly assigned labels (Fig. 3a). The *F*1 score achieved by the background model is approximately 50%, which is close to the random chance (mean *F*1 = 0.522), while the true model attained a significantly higher mean *F*1 score of 0.737. This indicates that PlantRNA-FM(+) has successfully learned the RNA features in the 5’ UTR sequences associated with translation.

**Fig. 3.**
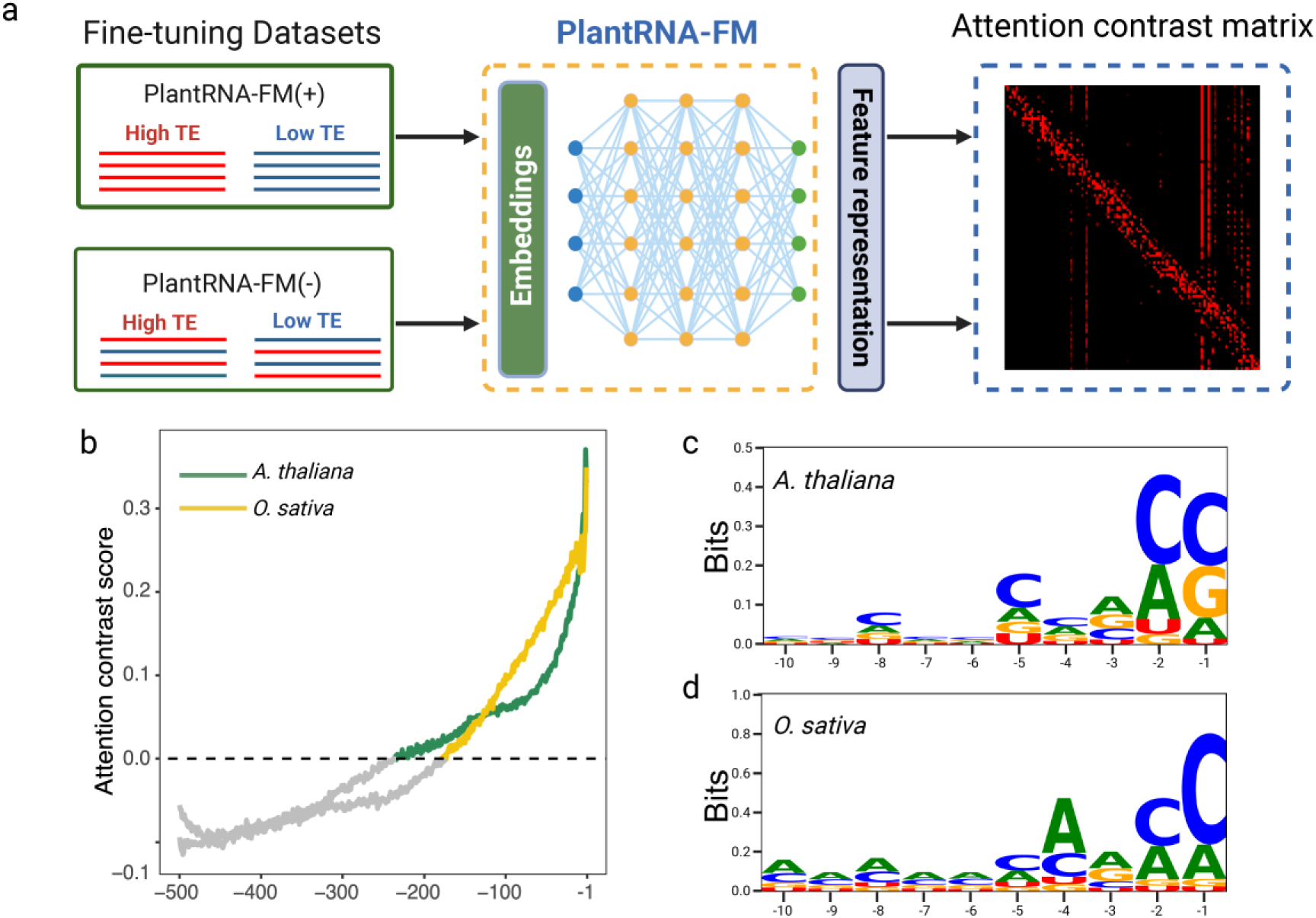
Our model interpretable framework reveals translation-associated RNA features. a, Schematic of the model interpretability approach. b, Transcriptome-wide attention contrast scores. The −1 position represents the first site upstream of the AUG. Different species are distinguished by colours. c, d, The information content of the 10 high-attention bases closest to the AUG start codon.

By subtracting the attention matrices of the background model from those of the true model, we obtained an attention contrast matrix that highlighted the significance of nucleotides in the 5’ UTR contributing to TE (Fig. 3a). Across the transcriptomes, we observed an increase in attention contrast scores as the position approached the AUG start codon in both *Arabidopsis* and rice (Fig. 3b). This result indicates that positions close to the start codon contribute the most to the TE values. By underlining the RNA sequence contents with high contrast attention score (identified by a z-score > 2.326), our PlantRNA-FM successfully identified the Kozak sequence motifs in both *Arabidopsis* and rice transcriptomes that are associated with TE (Fig. 3c, 3d). This result demonstrates that our PlantRNA-FM successfully identifies evolutionarily conserved RNA motifs that are important to translation (Fig. 3c, 3d).

### 1.4. PlantRNA-FM globally identifies the translation-associated RNA secondary structure motifs

Since RNA structure is the unique RNA feature incorporated in our PlantRNA-FM, we further identified the RNA secondary structure motifs important to translation through the model’s attention contrast matrix and an unsupervised hierarchical clustering strategy (Fig. 4a, **Methods**). Overall, we identified 112 RNA secondary structure motifs that are important to translation, including 63 low translation-associated and 49 high translation-associated RNA secondary structure motifs (Table S2). Notably, we identified low translation-associated RNA secondary structure motifs with high GC base pairs such as the RNA secondary structure motif with four GC base pairs in the stem (Fig. 4b). Interestingly, we also identified high translation-associated RNA structure motifs with a balanced ratio of GC and AU base pairs such as the RNA structure motif with four base pairs formed by two repeats of ACGU (Fig. 4c).

**Fig 4.**
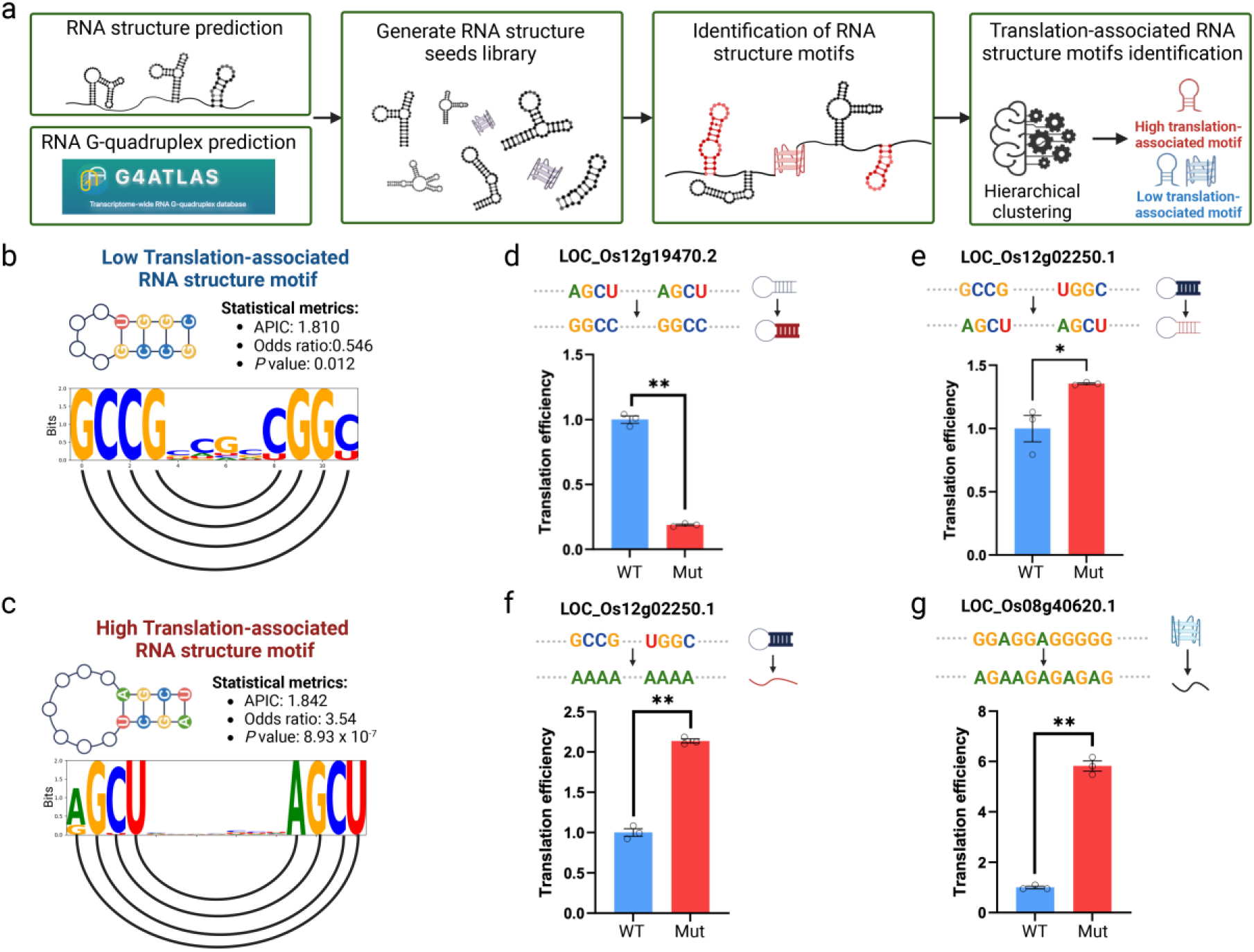
RNA structure motif identification approach reveals translation-associated RNA structure motifs. a, Overview of the RNA structure motif identification approach. RNA structures are predicted using RNAfold with a maximum length of 30 nucleotides to obtain RNA structure seeds. Predicted RNA G-quadruplexes were obtained from the G4Atlas database. b, c, Schematic diagram of high translation-associated RNA structure motifs and low translation-associated RNA structure motifs. Sequence logos show the information content of each nucleotide, with semicircles connecting paired bases. APIC stands for average positional information content. The *p-value* is derived from Fisher’s exact test. d,e,f,g, Experimental validation of high and low translation-associated RNA structure motifs and low translation-associated RNA G-quadruplex. The bar plot represents the translational efficiency of the original (WT) and RNA structure-mutated (Mut) constructs from the dual luciferase reporter assay in plants. It represents the change from high translation-associated RNA structure motifs to low translation-associated RNA structure motifs (d), the change from low translation-associated RNA structure motifs to high translation-associated RNA structure motifs (e), the complete disruption of low translation-associated RNA structure motifs (f), and the complete disruption of low translation-associated rG4 (g). * indicates *P* < 0.05, ** indicates *P* < 0.01, by Student’s t-test, n = 3, error bars indicate se.

To validate our identified RNA secondary structure motifs important to translation, we conducted experimental validation using the dual luciferase reporter assay in plants^12^. For the high translation-associated RNA secondary structure motif with four base pairs formed by two repeats of ACGU, we changed the two AU base pairs to the two GC base pairs, resulting in a significant decrease in TE with a reduction up to 5.3-fold (Fig. 4d). In contrast, when we exchanged the low translation-associated RNA secondary structure motif with four GC base pair in the stem for the high translation-associated RNA secondary structure motif with a balanced mix of GC and AU base pairs, we found a significant increase in TE (Fig. 4e). Notably, when we completely disrupted this low translation-associated RNA structure motif, resulting in complete single-strandedness, we observed an even greater enhancement of TE up to 2.1-fold (Fig. 4f). Our results demonstrate that PlantRNA-FM is capable of determining functional RNA secondary structure motifs in plants.

### 1.5. PlantRNA-FM globally identifies the translation-associated RNA tertiary structure motifs

RNA G-quadruplexes (rG4s) are one of the RNA tertiary structure motifs formed by the stacking of two or more G-quartets, composed of four guanines held together by both Watson-Crick and Hoogsteen hydrogen bonds^8,32,33^. Previous studies have demonstrated the important role of individual rG4s in repressing translation^34^. However, it is impossible to identify all the rG4 motifs important to translation from tens of thousands of rG4 motifs across the transcriptome. Therefore, we took advantage of our PlantRNA-FM to identify the translation-associated rG4s at the transcriptome-wide scale.

We first obtained all rG4 motifs in the 5’ UTRs from our G4Atlas database^33^. Subsequently, we identified all rG4 motifs associated with translation using our model’s attention contrast matrix across the transcriptome (**Methods**). Notably, we only identified rG4 motifs associated with low TE, particularly with both GGA and GGU repeat (Table S3). Therefore, our results indicate that rG4 serves as a translation repressor, which agrees with previous studies on individual rG4s^35–37^. To validate our identified translation-associated rG4 motifs, we conducted the experimental validation using dual luciferase reporter assay in plants^12^. We fused the 5’UTRs containing our identified rG4 motif and the corresponding disrupted rG4 motif with the luciferase reporter genes^12^. We then measured the corresponding TEs in plants and observed a significant increase of up to 5.8-fold in the disrupted rG4 motif compared to the TE in the native rG4 motif (Fig. 4g). These results indicate that our PlantRNA-FM is also capable of identifying functional RNA tertiary structure motifs such as translation-associated rG4 motifs throughout the transcriptome.

## 2. Discussion

In this study, we developed PlantRNA-FM, a high-performance and interpretable plant-specific RNA FM. PlantRNA-FM (Fig. 1) is designed for understanding RNA sequence and structure information rather than generation. This state-of-the-art model was specifically designed based on the extensive plant RNA information from 1,124 plant species, thereby capturing the remarkable diversity of plant RNA features. From the perspective of the dataset, we have incorporated RNA sequence information of all the RNAs from the transcriptomes across 1,124 plant species. We also incorporated the corresponding RNA annotation information. The integration of RNA structure information in our PlantRNA-FM achieves superior performance in RNA structure prediction tasks compared to other FMs (Fig. S1). Regarding the model architecture, we adopted a fine-grained tokenization method with single-nucleotide resolution. This contrasts with commonly used tokenization methods, such as byte pair encoding (BPE) and k-mers, which rely on frequency-based tokenization and may inadvertently fragment RNA structure motifs into arbitrary pieces. This strategy ensures the precise extraction and preservation of RNA structure motifs as coherent units throughout the pre-training process, thereby maintaining the integrity of crucial structure information. Additionally, PlantRNA-FM integrates rotational position embedding (RoPE), a technique that has proven effective in enhancing the modeling capabilities for long tokens in large FMs^38^. The implementation of RoPE leads to a approximately 30% reduction in the number of parameters in the embedding layer, consequently improving the efficiency of RNA tokenisation and modeling.

The superior performance of PlantRNA-FM can be further demonstrated in the plant-specific downstream tasks (Fig. 2a). Our PlantRNA-FM achieved the best *F1* scores of 0.974 and 0.958 for the genic region annotation in *Arabidopsis* and rice, while our PlantRNA-FM also achieved much better performance in predicting TE compared to other FMs (Fig. 2b, 2c). The outperformance of our PlantRNA-FM is likely due to the combination of both RNA sequence and structure information in our pre-training dataset, highlighting the importance of RNA structure, a key RNA feature, in regulating RNA biological processes.

Notably, we developed an interpretable framework for our PlantRNA-FM to explore the RNA features within the 5’ UTR that influence translation (Fig. 3a). Using the attention contrast matrices, we found that the nucleotides in the regions close to the start codon affect the translation the most, emphasizing the importance of positional information of functional RNA motifs (Fig. 3b). In contrast to conventional meta-gene analysis, our PlantRNA-FM is capable of providing positional information of RNA motifs across transcriptomes, which is critical for biological regulatory functions. Furthermore, the Kozak sequence, an evolutionary conserved translation-associated sequence motif across translation initiation sites was successfully identified in both *Arabidopsis* and rice using our PlantRNA-FM (Fig. 3c, 3d). This result successfully demonstrates the capability of our PlantRNA-FM in identifying the RNA sequence motifs important to translation across the transcriptomes. By using an unsupervised hierarchical clustering strategy to explore our attention contrast matrix, we further systematically identified RNA secondary and tertiary structure motifs that are functionally important to translation (Fig. 4a). Notably, we identified both high translation-associated and low translation-associated RNA secondary structure motifs where their differences are mainly in the strengths of the base pairs (Fig. 4b, 4c). This suggests that RNAs may adopt different RNA structure motifs with diverse folding strengths in regulating biological processes such as translation. In contrast to conventional meta-gene analysis, our PlantRNA-FM is capable of delivering a comprehensive understanding of functional RNA motifs such as the type of RNA motifs, the genic position of the RNA motifs, the positive or negative effects of the RNA motifs on their functions, and the exact contributions of the RNA motifs to their functions. For instance, high GC content in the 5’ UTR has been shown to be anti-correlated with translation efficiency^39–41^. However, these correlations are not able to facilitate understanding of which type of regulatory motifs with high GC content repress translation. Here, our PlantRNA-FM revealed diverse RNA structure motifs such as the RNA secondary structure motif with four GC base pairs in the stem and rG4s, serving as low translation-associated RNA motifs. This suggests the diversity of RNA regulatory motifs across the transcriptomes (Fig. 4b).

In summary, we have built the first interpretable RNA FM with both RNA sequence and structure information. Our PlantRNA-FM was pre-trained using 1,124 plant transcriptomes. We have demonstrated that our PlantRNA-FM is capable of identifying functional RNA motifs such as translation-associated sequence and structure motifs across the transcriptomes. Through our experimental validations, we have elucidated novel translation-associated RNA motifs in plants. Our FM model can be extended to explore functional RNA motifs in other kingdoms and investigate RNA motifs important for other biological functions such as RNA decay and maturation. Our PlantRNA-FM is poised to transform the way we determine RNA motifs for regulating gene expression, opening new horizons for programming RNA codes to facilitate crop improvements and RNA-based applications.

## 3. Methods

### 3.1. Pre-training datasets curation

The plant transcriptome data used for pre-training PlantRNA-FM was obtained from the one thousand plant transcriptomes project (1KP)^14^. Note that modeling genomic sequences differs significantly from natural language modeling. For instance, while RNA sequences are one-dimensional, they strictly follow biological genomic patterns and depend heavily on certain structural characteristics. In contrast, natural language models are more resilient and can tolerate linguistic errors such as typos and grammar mistakes. Thus, effective RNA sequence curation is crucial to minimize the impact of noisy data and enhance modeling performance. Specifically, our data curation protocol is as follows.

- **Sequence truncation and filtering**: We truncated RNA sequences exceeding 512 nucleotides to comply with the model’s maximum length capacity and filtered out sequences shorter than 20 nucleotides to eliminate noise, such as RNA fragment sequences.
- **RNA secondary structure annotation:** Given the significant impact of RNA secondary structures on sequence function, we annotated the local RNA structures of all RNA sequences using ViennaRNA (with parameters maxBPspan = 30)^25^.
- **Annotation of CDS and UTR sequences:** After obtaining the assembled transcripts and translated RNA regions from the dataset, we retrieve the CDS (translated RNA), 5’ UTR, and 3’ UTR sequences (upstream and downstream of the translated RNA).

### 3.2. Model architecture

In this study, we developed PlantRNA-FM, a specialised language model based on the transformer architecture (Fig. 1). PlantRNA-FM has 35 million parameters, including 12 transformer network layers, 24 attention heads, and an embedding dimension of 480. We applied layer normalisation and residual connections both before and after the encoder block. As our focus is on RNA understanding rather than generation, we only utilised the encoder component of the transformer architecture. PlantRNA-FM is capable of processing sequences up to 512 nucleotides in length, making it compatible with consumer-grade GPUs, such as the Nvidia RTX 4090, with a batch size of 16. The model was trained on four A100 GPUs over a period of three weeks, completing 3 epochs.

### 3.3. Pretraining strategies of PlantRNA-FM

To develop an RNA FM for exploiting all potential patterns within RNA sequences, we investigated the biological domain knowledge of RNA sequences and propose three self-supervised pre-training objectives to enhance the foundational model.

#### 3.3.1. Pretraining with Masked nucleotides modeling

Inspired by the concept of masked language modelling (MLM) in NLP, we introduced masked nucleotide modelling (MNM) for RNA sequences. This approach involves randomly masking a portion of nucleotides and leveraging the model itself to reconstruct these masked nucleotides. Note that the ability to accurately reconstruct masked nucleotides indicates that the model is empowered with the capability of understanding RNA sequence. MNM dynamically selects 20% of nucleotides for masking in each input sequence, as opposed to the fixed 15% masking used in the classic MLM objective designed for shorter natural language sentences. This increased masking ratio is chosen to enhance MNM’s modeling capability, considering that RNA sequences typically contain around one thousand bases. Specifically, 10% are replaced with a ‘ <mask>’ token, 5% with random nucleotides, and the remaining 5% are left as is. This approach, which aims for token classification, employs cross-entropy as the loss function to enhance the model’s predictive accuracy for masked or replaced nucleotides. The loss function *L*_*MLM*_(*θ*) for MLM is defined as follows:

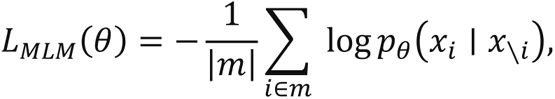

where *θ* and *m* are the parameter set inside the FM and the number of masked nucleotides. *p*_*θ*_(*x*_*i*_ ∣ *x*_∖*i*_) indicates the probability of predicting the masked nucleotide *x*_*i*_ based on its context (*x*_∖*i*_).

#### 3.3.2. Pretraining with RNA Structure Prediction

We hypothesise that effectively aligning RNA sequences with their corresponding secondary structures is important during the pre-training phase. In practice, we annotated the secondary structures within the 1KP dataset, which comprises 50 billion nucleotides. This establishes a robust foundation for our model to recognise the critical role of secondary structures. Based on these annotated data, we utilized cross-entropy as the loss function to predict the RNA secondary structure:

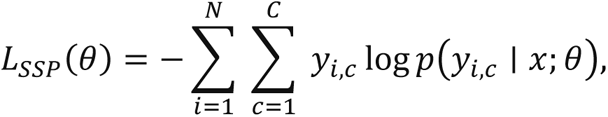

where *N* is the length of the RNA sequence, i.e., the total number of nucleotides in the sequence; *C* denotes the number of prediction for each nucleotide (e.g., ‘(’, ‘)’, ‘.’); *y*_*i*,*c*_ is the prediction of the *i*-th nucleotide *c*, and *p*(*y*_*i*,*c*_ ∣ *x*; *θ*) is the probability predicted by the model parameterised by *θ*. *L*_*SSP*_ (*θ*) is the loss function that quantifies the discrepancy between the model’s predicted probabilities for each nucleotide’s secondary structure and the actual structure, with the aim of minimising this loss to improve the model’s accuracy in secondary structure prediction.

#### 3.3.3. Pretraining with RNA annotation prediction

RNA sequences exhibit significant variation across different regions, each serving distinct functions within an organism. Beyond the two aforementioned training objectives, the third one focuses on classifying regions within RNA sequences. The loss function is as follows:

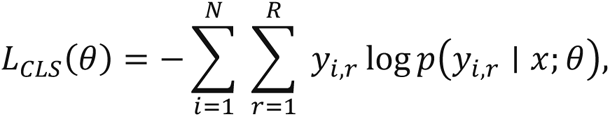

where *N* is the length of the RNA sequence, i.e., the total number of nucleotides or segments considered for classification. *R* represents the number of region categories we are classifying, including CDS, 3’ UTR, and 5’ UTR. *y*_*i*,*r*_ is the prediction of the *i*-th nucleotide *r*. *p*(*y*_*i*,*r*_ ∣ *x*; *θ*) is the probability predicted by the model, with parameters *θ*, for the *i* -th nucleotide given the RNA sequence *x*. *L*_*CLS*_(*θ*) is the cross-entropy loss function aimed at training the model to identify different regions.

### 3.4. Fine-tuning of downstream tasks

After the pre-training phase, our FM can be fine-tuned to adapt to various downstream tasks. The fine-tuning phase consists of three steps. First, we gathered an annotated dataset specific to each downstream task, which consists of sequences and their corresponding labels. Note that we pre-sliced any sequences that exceed the model’s maximum length, to ensure compatibility. Next, using the pre-trained FM as a starting point, we adapted the output layer to accommodate the requirements of RNA modelling tasks, which may include outputting sequences, labels, or scalar values. Finally, the training and inference processes are tailored to the demands of each downstream task by selecting task-specific optimisers, loss functions, and tuning hyperparameters to achieve optimal performance. The source code for our training and inference can be found in our repository.

### 3.5. Polysome profiling mapping and data processing

Raw polysome profiling sequencing data for *A. thaliana* were obtained from published research^12^. For rice, we performed polysome-seq using the same protocol as *Arabidopsis*^8^. The genomes and annotation files of *O. sativa* and *A. thaliana* were obtained from Phytozome v13 with version of *Oryza sativa* v7.0 and TAIR10^44^. After extracting the transcriptome sequence through the reference genome and annotation files, clean polysome profiling and RNA-Seq reads were mapped to the reference transcriptome using HISAT2 and followed by library normalisation and quantification using DESeq2^45,46^. Next, genes with an RPKM of less than 1 were removed, and the TE of each gene was calculated by dividing the polysome-associated RNA levels (polysome profiling RNA-seq) by the corresponding RNA levels (RNA-Seq)^12^. Subsequently, the dataset was classified as high or low TE, using the mean plus or minus the standard deviation as a threshold, and were respectively assigned the labels 1 and 0 for high and low TE.

### 3.6. RNA structure motif identification approach

#### 3.6.1. Extraction of the attention contrast matrix

To facilitate better model interpretation, we created two additional models. One is the true model, denoted as PlantRNA-FM(+), trained using the real TE labels, while the other one is the background model, PlantRNA-FM(-), altered using the same dataset but with randomly assigned labels. Specifically, we fine-tuned the pre-trained PlantRNA-FM (+) and (-) on each dataset for 100 epochs, using regular hyperparameter settings. To avoid overfitting, we employed an early stopping strategy to terminate the fine-tuning process when the best *F1* score remained unchanged for 30 epochs. Once the fine-tuning was completed, we used the fine-tuned models to predict each dataset and derive the raw attention score matrices corresponding to each RNA sequence. Since the raw attention score matrices are five-dimensional, we reshaped them through average-based downsampling to generate attention contrast matrices. Finally, we subtracted the attention contrast matrices of PlantRNA-FM (+) from those of PlantRNA-FM (-). Furthermore, we padded any negative values in the attention contrast matrices with zeros for better visualisation.

#### 3.6.2. Generation of the RNA structure motif seed library

To identify RNA structure motifs, we first generate a library of that contains RNA structure motif seeds derived from RNA sequences across the transcriptomes. In this work, we apply the Zuker algorithm from the Vienna RNA package to obtain all suboptimal RNA structure foldings for each RNA in our dataset^35,36^. We restrict the length of the RNA structure motifs to a maximum of thirty^48^. The folded RNA structures are then annotated using “bpRNA”. Subsequently, all RNA structure motifs are extracted to generate a seed library of RNA structure motifs for the plant transcriptomes^26^. In order to obtain reliable RNA structure motifs, we set the range of RNA structure stems from 4 to 7, and the loop length from 4 to 9.

#### 3.6.3. Identification of translation-associated RNA secondary structure motifs

From the previous step, we obtained all potential foldings of the RNA structure motif in the 5’ UTR and aligned them with the attention contrast matrix. For each RNA structure motif, we evaluated it using a paired *t*-test to obtain a *p*-value. Then, we corrected the obtained *p*-value using the Benjamini-Hochberg (BH) method. RNA structure motifs with *p*-values less than 0.01 were considered significant and extracted as the high-attention RNA structure motifs. Then we extracted their corresponding RNA sequence and converted them into numerical matrices using the one-hot encoding method. Subsequently, we applied an unsupervised hierarchical clustering strategy to classify the nucleotides corresponding to the positions of the RNA structure pairs into 2 to 100 clusters^49^. For each cluster containing a minimum of 30 high-attention RNA structure motifs, the significance was assessed using Fisher’s exact test. RNA motifs with an odds ratio over 1 and a *p*-value below 0.05 were identified as high translation-associated motifs. On the contrary, those with an odds ratio less than 1 and a *p*-value below 0.05 were associated with low TE. Additionally, we calculated the mean information content of all bases, defined as the “Average Positional Information Content” (APIC). RNA motifs with an APIC below 1.5 were excluded from further analysis.

#### 3.6.4. Identification of translation-associated rG4s

We obtained all potential rG4 in rice from our G4Atlas database^33^. Next, we aligned the rG4 sequences with the corresponding attention contrast matrix and employed the paired *t*-test to assess the statistical significance. For each length of rG4, we adjusted its *p*-value using the Benjamini-Hochberg (BH) correction method and selected rG4s with a *p*-value less than 0.01 as the high attention rG4s.

## 4. Data availability

The polysome-seq sequence data of *A. thaliana* was obtained from the Sequence Read Archive (SRA) (https://www.ncbi.nlm.nih.gov/sra) under BioProject ID number PRJNA762705^8^. The raw sequence data of *O. sativa* has been deposited in the Sequence Read Archive (SRA) (https://www.ncbi.nlm.nih.gov/sra) under BioProject ID number PRJNA1112739.

## 5. Code availability

The source code of this study is freely available at Huggingface (https://huggingface.co/yangheng/PlantRNA-FM).

## References

1. Piao, M., Sun, L. & Zhang, Q. C. RNA Regulations and Functions Decoded by Transcriptome-wide RNA Structure Probing. Genomics, Proteomics & Bioinformatics 15, 267–278 (2017).

2. Komatsu, K. R. et al. RNA structure-wide discovery of functional interactions with multiplexed RNA motif library. Nature Communications 11, 6275 (2020).

3. Espah Borujeni, A., Channarasappa, A. S. & Salis, H. M. Translation rate is controlled by coupled trade-offs between site accessibility, selective RNA unfolding and sliding at upstream standby sites. Nucleic Acids Research 42, 2646–2659 (2014).

4. Gorochowski, T. E., Ignatova, Z., Bovenberg, R. A. L. & Roubos, J. A. Trade-offs between tRNA abundance and mRNA secondary structure support smoothing of translation elongation rate. Nucleic Acids Research 43, 3022–3032 (2015).

5. Mortimer, S. A., Kidwell, M. A. & Doudna, J. A. Insights into RNA structure and function from genome-wide studies. Nat Rev Genet 15, 469–479 (2014).

6. Kozak, M. An analysis of vertebrate mRNA sequences: intimations of translational control. Journal of Cell Biology 115, 887–903 (1991).

7. Ding, Y. et al. In vivo genome-wide profiling of RNA secondary structure reveals novel regulatory features. Nature 505, 696–700 (2014).

8. Yang, X. et al. RNA G-quadruplex structure contributes to cold adaptation in plants. Nat Commun 13, 6224 (2022).

9. Xu, B. et al. Recent advances in RNA structurome. Sci. China Life Sci. 65, 1285–1324 (2022).

10. Yang, M., et al. Intact RNA structurome reveals mRNA structure-mediated regulation of miRNA cleavage in vivo. bioRxiv 2019.12.21.885699 (2020) doi:10/ghccqf.

11. Yang, M. et al. In vivo single-molecule analysis reveals COOLAIR RNA structural diversity. Nature 1–6 (2022) doi:10.1038/s41586-022-05135-9.

12. Xiaofei Yang & Haopeng Yu. Wheat in vivo RNA structure landscape reveals a prevalent role of RNA structure in modulating translational subgenome expression asymmetry. 26 (2021).

13. Deng, H. et al. Rice In Vivo RNA Structurome Reveals RNA Secondary Structure Conservation and Divergence in Plants. Molecular Plant 11, 607–622 (2018).

14. One Thousand Plant Transcriptomes Initiative. One thousand plant transcriptomes and the phylogenomics of green plants. Nature 574, 679–685 (2019).

15. Cao, J. et al. High-throughput 5′ UTR engineering for enhanced protein production in non-viral gene therapies. Nat Commun 12, 4138 (2021).

16. Consens, M. E., et al. To Transformers and Beyond: Large Language Models for the Genome. Preprint at 10.48550/arXiv.2311.07621 (2023).

17. Zhou, Z., et al. DNABERT-2: Efficient Foundation Model and Benchmark For Multi-Species Genome. Preprint at 10.48550/arXiv.2306.15006 (2023).

18. Dalla-Torre, H. et al. The Nucleotide Transformer: Building and Evaluating Robust Foundation Models for Human Genomics. 2023.01.11.523679 Preprint at 10.1101/2023.01.11.523679 (2023).

19. Lin, Z. et al. Evolutionary-scale prediction of atomic-level protein structure with a language model. Science 379, 1123–1130 (2023).

20. Chu, Y. et al. A 5’ UTR Language Model for Decoding Untranslated Regions of mRNA and Function Predictions. 2023.10.11.561938 Preprint at 10.1101/2023.10.11.561938 (2023).

21. Hallee, L., Rafailidis, N. & Gleghorn, J. P. cdsBERT - Extending Protein Language Models with Codon Awareness. bioRxiv 2023.09.15.558027 (2023) doi:10.1101/2023.09.15.558027.

22. Chen, K. et al. Self-supervised learning on millions of primary RNA sequences from 72 vertebrates improves sequence-based RNA splicing prediction. Briefings in Bioinformatics 25, bbae163 (2024).

23. Yang, X., Yang, M., Deng, H. & Ding, Y. New Era of Studying RNA Secondary Structure and Its Influence on Gene Regulation in Plants. Front. Plant Sci. 9, (2018).

24. Zhang, H., Chung, B. Y.-W., Wang, Z. & Ding, Y. Editorial: Plant RNA structure. Front. Plant Sci. 14, (2023).

25. Lorenz, R. et al. ViennaRNA Package 2.0. Algorithms Mol Biol 6, 26 (2011).

26. Danaee, P. et al. bpRNA: large-scale automated annotation and analysis of RNA secondary structure. Nucleic Acids Research 46, 5381–5394 (2018).

27. Sloma, M. F. & Mathews, D. H. Exact calculation of loop formation probability identifies folding motifs in RNA secondary structures. RNA 22, 1808–1818 (2016).

28. Tan, Z., Fu, Y., Sharma, G. & Mathews, D. H. TurboFold II: RNA structural alignment and secondary structure prediction informed by multiple homologs. Nucleic Acids Research 45, 11570–11581 (2017).

29. Hardy, E. C. & Balcerowicz, M. Untranslated yet indispensable—UTRs act as key regulators in the environmental control of gene expression. Journal of Experimental Botany erae073 (2024) doi:10.1093/jxb/erae073.

30. Dever, T. E., Ivanov, I. P. & Hinnebusch, A. G. Translational regulation by uORFs and start codon selection stringency. Genes Dev. 37, 474–489 (2023).

31. Evfratov, S. A. et al. Application of sorting and next generation sequencing to study 5΄-UTR influence on translation efficiency in Escherichia coli. Nucleic Acids Research 45, 3487–3502 (2017).

32. Lyu, K., Chow, E. Y.-C., Mou, X., Chan, T.-F. & Kwok, C. K. RNA G-quadruplexes (rG4s): genomics and biological functions. Nucleic Acids Research 49, 5426–5450 (2021).

33. Yu, H., Qi, Y., Yang, B., Yang, X. & Ding, Y. G4Atlas: a comprehensive transcriptome-wide G-quadruplex database. Nucleic Acids Research 51, D126–D134 (2023).

34. Song, J., Perreault, J.-P., Topisirovic, I. & Richard, S. RNA G-quadruplexes and their potential regulatory roles in translation. Translation 4, e1244031 (2016).

35. Kumari, S., Bugaut, A., Huppert, J. L. & Balasubramanian, S. An RNA G-quadruplex in the 5′ UTR of the NRAS proto-oncogene modulates translation. Nat Chem Biol 3, 218– 221 (2007).

36. Beaudoin, J.-D. & Perreault, J.-P. 5′-UTR G-quadruplex structures acting as translational repressors. Nucleic Acids Research 38, 7022–7036 (2010).

37. Jia, L. et al. Decoding mRNA translatability and stability from the 5′ UTR. Nat Struct Mol Biol 27, 814–821 (2020).

38. Su, J. et al. RoFormer: Enhanced transformer with Rotary Position Embedding. Neurocomputing 568, 127063 (2024).

39. Araujo, P. R. et al. Before It Gets Started: Regulating Translation at the 5′ UTR. International Journal of Genomics 2012, e475731 (2012).

40. Leppek, K., Das, R. & Barna, M. Functional 5′ UTR mRNA structures in eukaryotic translation regulation and how to find them. Nat Rev Mol Cell Biol 19, 158–174 (2018).

41. van der Velden, A. W. & Thomas, A. A. M. The role of the 5′ untranslated region of an mRNA in translation regulation during development. The International Journal of Biochemistry & Cell Biology 31, 87–106 (1999).

42. Jumper, J. et al. Highly accurate protein structure prediction with AlphaFold. Nature 596, 583–589 (2021).

43. Verkuil, R. et al. Language models generalize beyond natural proteins. 2022.12.21.521521 Preprint at 10.1101/2022.12.21.521521 (2022).

44. Goodstein, D. M. et al. Phytozome: a comparative platform for green plant genomics. Nucleic Acids Research 40, D1178–D1186 (2012).

45. Kim, D., Langmead, B. & Salzberg, S. L. HISAT: a fast spliced aligner with low memory requirements. Nat Methods 12, 357–360 (2015).

46. Love, M. I., Huber, W. & Anders, S. Moderated Estimation of Fold Change and Dispersion for RNA-Seq Data with DESeq2. http://biorxiv.org/lookup/doi/10.1101/002832 (2014) doi:10.1101/002832.

47. Zuker, M. On Finding All Suboptimal Foldings of an RNA Molecule. Science 244, 48–52 (1989).

48. Fish, L. et al. A prometastatic splicing program regulated by SNRPA1 interactions with structured RNA elements. Science 372, eabc7531 (2021).

49. Steinbach, M., Karypis, G. & Kumar, V. A Comparison of Document Clustering Techniques. (2000).

