## Supplementary material for "PlantRNA-FM: An Interpretable RNA Foundation Model for Exploration Functional RNA Motifs in Plants": Fig. S1

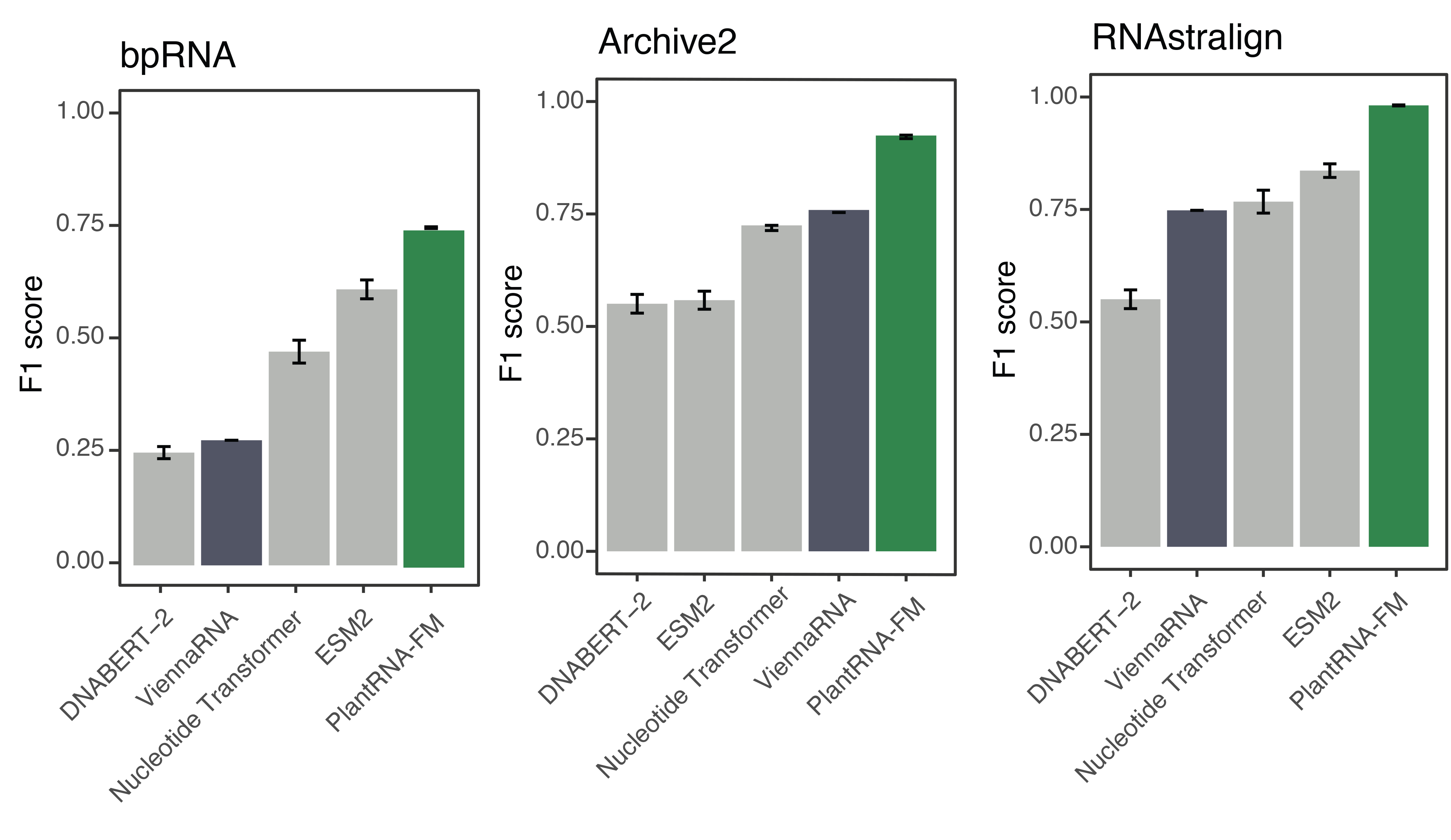

**Fig. S1. Comparison of the model performance of different pre-trained models on the RNA secondary structure prediction task.** Three classic RNA secondary structure datasets were used to evaluate the performance of different pre-trained models. The most widely used ViennaFold software was also considered and marked in dark colour. PlantRNA-FM is highlighted in green.
