## Supplementary material for "PlantRNA-FM: An Interpretable RNA Foundation Model for Exploration Functional RNA Motifs in Plants": Table S1

Table S1. Comparison of F1 scores of different pre-trained models on RNA structure prediction task

| Datasets | PlantRNA-FM | PlantRNA-FM-RNA-Only | ESM2 | DNABERT-2 | Nucleotide Transformer | ViennaRNA |
| --- | --- | --- | --- | --- | --- | --- |
| bpRNA-1m | 0.750±0.002 | 0.694±0.001 | 0.614±0.017 | 0.251±0.011 | 0.475±0.0208 | 0.278 |
| ArchiveII | 0.924±0.002 | 0.912±0.001 | 0.558±0.016 | 0.550±0.021 | 0.725±0.0047 | 0.759 |
| RNAstralign | 0.981±0.001 | 0.976±0.001 | 0.836±0.012 | 0.616±0.004 | 0.767±0.0208 | 0.748 |
