## Supplementary material for "PlantRNA-FM: An Interpretable RNA Foundation Model for Exploration Functional RNA Motifs in Plants": Table S2

**Table S2. Identified RNA structure motifs associated translation efficiency.**

| **Label** | **RNA Structure** | **Consensus_base** | **OddsRatio** | **P value** | **APIC** |
| --- | --- | --- | --- | --- | --- |
| low TE-related | ((((......)))) | GGGG......TCCC | 0.165 | 3.081E-05 | 1.679 |
| low TE-related | ((((......)))) | GGGG......TCCC | 0.312 | 3.694E-05 | 1.522 |
| low TE-related | ((((.........)))) | CGCC.........GGCG | 0.307 | 7.556E-05 | 2.000 |
| low TE-related | ((((......)))) | CGCC......GGCG | 0.316 | 1.210E-04 | 2.000 |
| low TE-related | ((((......)))) | CGCC......GGCG | 0.451 | 2.689E-04 | 1.625 |
| low TE-related | ((((......)))) | CGTC......GGCG | 0.535 | 3.750E-04 | 1.567 |
| low TE-related | ((((.......)))) | CCCC.......GGGG | 0.276 | 3.809E-04 | 1.771 |
| low TE-related | ((((....)))) | CCGC....GCGG | 0.343 | 4.618E-04 | 2.000 |
| low TE-related | ((((.......)))) | AGGG.......CCCT | 0.368 | 9.016E-04 | 1.529 |
| low TE-related | ((((.......)))) | CCGC.......GCGG | 0.381 | 1.127E-03 | 2.000 |
| low TE-related | ((((........)))) | GCTC........GGGC | 0.261 | 1.346E-03 | 1.636 |
| low TE-related | (((((....))))) | CGCCG....CGGCG | 0.362 | 2.037E-03 | 1.624 |
| low TE-related | ((((.........)))) | CTCC.........GGAG | 0.494 | 2.408E-03 | 1.649 |
| low TE-related | ((((....)))) | TCCC....GGGG | 0.388 | 2.526E-03 | 1.540 |
| low TE-related | ((((......)))) | GGGG......CCCC | 0.388 | 2.529E-03 | 1.667 |
| low TE-related | ((((.......)))) | GGGG.......TCCC | 0.322 | 2.585E-03 | 1.663 |
| low TE-related | ((((.......)))) | GGCG.......CGCC | 0.372 | 3.065E-03 | 1.894 |
| low TE-related | ((((........)))) | GGGG........CTCC | 0.372 | 3.065E-03 | 1.686 |
| low TE-related | ((((......)))) | TTCC......GGAG | 0.461 | 3.261E-03 | 1.669 |
| low TE-related | ((((.....)))) | GGAG.....TTCC | 0.453 | 3.529E-03 | 1.584 |
| low TE-related | ((((.....)))) | TTCC.....GGAG | 0.453 | 3.529E-03 | 1.622 |
| low TE-related | ((((........)))) | TCCT........GGGA | 0.496 | 5.521E-03 | 1.541 |
| low TE-related | ((((........)))) | TGCT........GGCG | 0.378 | 6.643E-03 | 1.720 |
| low TE-related | ((((.......)))) | GCCG.......CGGC | 0.494 | 7.301E-03 | 2.000 |
| low TE-related | ((((.........)))) | CGCC.........GGCG | 0.597 | 7.403E-03 | 1.605 |
| low TE-related | (((((....))))) | CGTCG....CGGCG | 0.416 | 8.300E-03 | 1.539 |
| low TE-related | ((((....)))) | CCGC....GCGG | 0.543 | 9.064E-03 | 1.813 |
| low TE-related | ((((....)))) | GGAG....CTCC | 0.456 | 9.117E-03 | 1.664 |
| low TE-related | ((((........)))) | CGCC........GGCG | 0.477 | 9.892E-03 | 1.802 |
| low TE-related | ((((........)))) | CCCC........GGGG | 0.515 | 1.127E-02 | 1.573 |
| low TE-related | ((((....)))) | GCCG....CGGC | 0.597 | 1.145E-02 | 1.842 |
| low TE-related | ((((........)))) | GGCG........CGCC | 0.546 | 1.247E-02 | 1.810 |
| low TE-related | ((((........)))) | GGGG........TTCC | 0.510 | 1.249E-02 | 1.547 |
| low TE-related | (((((.......))))) | CGTCG.......CGGCG | 0.537 | 1.405E-02 | 1.624 |
| low TE-related | ((((.........)))) | GCCG.........CGGC | 0.612 | 1.457E-02 | 1.685 |
| low TE-related | ((((........)))) | TCCT........AGGA | 0.429 | 1.464E-02 | 1.706 |
| low TE-related | ((((....)))) | GGCG....CGCC | 0.483 | 1.482E-02 | 1.768 |
| low TE-related | ((((......)))) | CCTT......GAGG | 0.483 | 1.483E-02 | 1.557 |
| low TE-related | ((((......)))) | TCTC......GGGA | 0.411 | 1.501E-02 | 1.751 |
| low TE-related | ((((.......)))) | TCGC.......GCGG | 0.617 | 1.550E-02 | 1.716 |
| low TE-related | ((((.....)))) | CGCC.....GGCG | 0.599 | 1.669E-02 | 1.584 |
| low TE-related | ((((....)))) | GAGA....TCTC | 0.494 | 1.681E-02 | 1.684 |
| low TE-related | ((((.......)))) | CTCC.......GGAG | 0.409 | 1.859E-02 | 1.840 |
| low TE-related | ((((....)))) | GCGG....CCGC | 0.505 | 1.896E-02 | 1.605 |
| low TE-related | ((((.......)))) | CCTT.......GGGG | 0.477 | 2.016E-02 | 1.582 |
| low TE-related | ((((.........)))) | CTCC.........GGAG | 0.547 | 2.545E-02 | 1.795 |
| low TE-related | ((((........)))) | CCGC........GCGG | 0.548 | 2.558E-02 | 1.893 |
| low TE-related | ((((....)))) | GAGG....CCTC | 0.465 | 3.039E-02 | 1.671 |
| low TE-related | ((((.....)))) | GCGG.....TCGC | 0.594 | 3.147E-02 | 1.555 |
| low TE-related | ((((.........)))) | CCCT.........AGGG | 0.508 | 3.181E-02 | 1.588 |
| low TE-related | ((((.....)))) | AGGG.....CTCT | 0.510 | 3.212E-02 | 1.632 |
| low TE-related | ((((.........)))) | GGGG.........CCCT | 0.449 | 3.239E-02 | 1.657 |
| low TE-related | ((((.........)))) | GGCG.........CGCC | 0.449 | 3.239E-02 | 2.000 |
| low TE-related | ((((....)))) | CCCT....AGGG | 0.521 | 3.557E-02 | 1.501 |
| low TE-related | ((((.......)))) | GGCG.......CGCC | 0.635 | 3.957E-02 | 1.538 |
| low TE-related | ((((.....)))) | CTCT.....AGGG | 0.486 | 4.321E-02 | 1.638 |
| low TE-related | ((((......)))) | GCTC......GAGC | 0.544 | 4.332E-02 | 1.567 |
| low TE-related | ((((......)))) | CGAG......CTCG | 0.544 | 4.332E-02 | 1.617 |
| low TE-related | ((((......)))) | CTCT......GGAG | 0.544 | 4.332E-02 | 1.602 |
| low TE-related | (((((......))))) | TCGTC......GGCGG | 0.547 | 4.364E-02 | 1.570 |
| low TE-related | ((((....)))) | GCCG....CGGC | 0.619 | 4.598E-02 | 2.000 |
| low TE-related | ((((......)))) | GGGG......TTCC | 0.472 | 4.715E-02 | 1.758 |
| low TE-related | ((((.........)))) | GGAG.........CTCC | 0.539 | 4.865E-02 | 1.781 |
| high TE-related | ((((.....)))) | AGCT.....AGCT | 4.668 | 1.423E-09 | 1.982 |
| high TE-related | ((((.....)))) | AGCT.....AGCT | 2.952 | 2.849E-08 | 1.642 |
| high TE-related | ((((.......)))) | AGCT.......AGCT | 3.541 | 8.939E-07 | 1.886 |
| high TE-related | ((((....)))) | AGCT....AGCT | 2.713 | 5.708E-06 | 2.000 |
| high TE-related | ((((......)))) | AGCT......AGCT | 2.983 | 8.236E-06 | 2.000 |
| high TE-related | ((((.........)))) | AGCT.........AGCT | 3.569 | 1.602E-05 | 2.000 |
| high TE-related | ((((....)))) | AGCT....AGCT | 2.214 | 2.435E-05 | 1.853 |
| high TE-related | ((((.........)))) | AGCT.........AGCT | 2.855 | 4.682E-05 | 1.847 |
| high TE-related | ((((........)))) | TAGC........GCTA | 2.494 | 1.940E-04 | 1.528 |
| high TE-related | ((((.....)))) | TAGC.....GCTA | 4.815 | 3.054E-04 | 1.802 |
| high TE-related | ((((........)))) | TAGC........GCTA | 3.262 | 3.868E-04 | 1.721 |
| high TE-related | (((((.....))))) | TAGCT.....AGCTA | 3.566 | 4.557E-04 | 1.540 |
| high TE-related | (((((.......))))) | AGCTA.......TAGCT | 4.194 | 4.837E-04 | 1.720 |
| high TE-related | ((((....)))) | GCTA....TAGC | 2.432 | 9.095E-04 | 1.829 |
| high TE-related | (((((....))))) | AGCTA....TAGCT | 3.162 | 1.553E-03 | 1.524 |
| high TE-related | ((((....)))) | GATC....GATC | 1.873 | 2.885E-03 | 1.814 |
| high TE-related | ((((....)))) | TCGA....TCGA | 2.126 | 2.984E-03 | 1.973 |
| high TE-related | ((((......)))) | CGAT......ATCG | 2.404 | 3.004E-03 | 1.943 |
| high TE-related | ((((....)))) | TAGC....GCTA | 2.833 | 3.075E-03 | 1.779 |
| high TE-related | ((((....)))) | GCTA....TAGC | 2.443 | 4.191E-03 | 2.000 |
| high TE-related | ((((......)))) | ATCG......CGAT | 2.269 | 7.260E-03 | 1.801 |
| high TE-related | (((((....))))) | CGATC....GATCG | 2.509 | 8.301E-03 | 1.891 |
| high TE-related | ((((.....)))) | CTAG.....CTAG | 2.794 | 8.955E-03 | 1.726 |
| high TE-related | ((((.....)))) | CTAG.....CTAG | 2.211 | 1.016E-02 | 1.516 |
| high TE-related | ((((.....)))) | GCTG.....TAGC | 1.965 | 1.030E-02 | 1.526 |
| high TE-related | (((((.....))))) | AGCTA.....TAGCT | 2.964 | 1.031E-02 | 1.597 |
| high TE-related | ((((......)))) | TTGC......GCAG | 2.420 | 1.205E-02 | 1.772 |
| high TE-related | (((((....))))) | TAGCT....AGCTA | 2.156 | 1.408E-02 | 1.649 |
| high TE-related | ((((........)))) | GCTA........TAGC | 2.146 | 1.424E-02 | 1.685 |
| high TE-related | ((((........)))) | AGCT........AGCT | 1.746 | 1.437E-02 | 1.836 |
| high TE-related | ((((....)))) | AAGC....GCTT | 2.289 | 1.438E-02 | 1.640 |
| high TE-related | ((((.........)))) | GCAG.........CTGC | 2.174 | 1.701E-02 | 1.680 |
| high TE-related | (((((......))))) | TAGCT......AGCTA | 2.277 | 1.721E-02 | 1.600 |
| high TE-related | ((((.....)))) | GATC.....GATC | 1.958 | 1.772E-02 | 1.687 |
| high TE-related | ((((......)))) | GCTA......TAGC | 2.218 | 2.034E-02 | 2.000 |
| high TE-related | ((((......)))) | GATC......GATC | 2.319 | 2.053E-02 | 1.865 |
| high TE-related | (((((....))))) | GCTAG....CTAGC | 2.012 | 2.253E-02 | 1.598 |
| high TE-related | ((((......)))) | TAGC......GCTA | 1.908 | 2.395E-02 | 1.805 |
| high TE-related | (((((....))))) | ATCGA....TCGAT | 1.994 | 2.585E-02 | 1.573 |
| high TE-related | ((((.......)))) | TAGC.......GCTA | 1.985 | 2.587E-02 | 1.830 |
| high TE-related | ((((.......)))) | CTGC.......GCAG | 1.682 | 3.483E-02 | 1.612 |
| high TE-related | ((((......)))) | GCTA......TAGC | 1.717 | 3.841E-02 | 1.668 |
| high TE-related | ((((....)))) | CTCG....CGAG | 1.868 | 3.879E-02 | 1.763 |
| high TE-related | ((((....)))) | GCTC....GAGC | 2.361 | 4.194E-02 | 1.775 |
| high TE-related | ((((....)))) | CTAG....CTAG | 1.625 | 4.292E-02 | 1.948 |
| high TE-related | ((((.......)))) | CTGC.......GCAG | 1.824 | 4.685E-02 | 1.766 |
| high TE-related | ((((.......)))) | GCTA.......TAGC | 1.824 | 4.685E-02 | 1.845 |
| high TE-related | ((((......)))) | AAGC......GATT | 2.030 | 4.795E-02 | 1.516 |
| high TE-related | ((((.........)))) | GAGC.........GCTC | 2.189 | 4.971E-02 | 1.632 |
