## Supplementary material for "PlantRNA-FM: An Interpretable RNA Foundation Model for Exploration Functional RNA Motifs in Plants": Table S3

Table S3. Identified RNA G-quadruplex associated translation efficiency.

| **RG4 sequence** | **Label** | **Odds ratio** | **P value** |
| --- | --- | --- | --- |
| GGAGGAGGAGG | Low TE-related | 0.421 | 8.68E-11 |
| GGGGGGGGGGG | Low TE-related | 0.404 | 1.05E-08 |
| GGGGGGGGGGGG | Low TE-related | 0.409 | 1.01E-07 |
| GGAGGAGGAGGAGG | Low TE-related | 0.429 | 2.86E-06 |
| GGGAGGAGGAGG | Low TE-related | 0.303 | 4.39E-06 |
| GGAGGAGGAGGG | Low TE-related | 0.303 | 4.39E-06 |
| GGGGGGGGGGGGG | Low TE-related | 0.439 | 4.78E-06 |
| GGGGGGGGGGGGGG | Low TE-related | 0.424 | 1.11E-05 |
| GGGGGGGGGGGGGGG | Low TE-related | 0.451 | 2.68E-04 |
| GGGGGAGGAGG | Low TE-related | 0.181 | 5.09E-04 |
| GGAGGAGGGGG | Low TE-related | 0.181 | 5.09E-04 |
| GGAGGAGGAGGAGGAGG | Low TE-related | 0.429 | 1.36E-03 |
| GGAGGAGGAGGAGGG | Low TE-related | 0.402 | 3.06E-03 |
| GGGAGGAGGAGGAGG | Low TE-related | 0.402 | 3.06E-03 |
| GGAGGAGGGGGG | Low TE-related | 0.086 | 3.91E-03 |
| GGGGGGAGGAGG | Low TE-related | 0.086 | 3.91E-03 |
| GGGGAGGAGGAGG | Low TE-related | 0.358 | 5.26E-03 |
| GGAGGAGGAGGGG | Low TE-related | 0.358 | 5.26E-03 |
| GGGGGGGGGGGGGGGG | Low TE-related | 0.507 | 7.55E-03 |
| GGAGGAGGAGGAGGAGGG | Low TE-related | 0.368 | 9.96E-03 |
| GGGAGGAGGAGGAGGAGG | Low TE-related | 0.368 | 9.96E-03 |
